# Editing of the Human TRIM5 Gene to Introduce Mutations with the Potential to Inhibit HIV-1

**DOI:** 10.1101/185298

**Authors:** Caroline Dufour, Alix Claudel, Nicolas Joubarne, Natacha Merindol, Tara Maisonnet, Mélodie B. Plourde, Lionel Berthoux

## Abstract

The type I interferon (IFN-I)-inducible human restriction factor TRIM5α inhibits the infection of human cells by specific nonhuman retroviruses, such as N-MLV and EIAV, but does not generally target HIV-1. However, the introduction of two aminoacid substitutions, R332G and R355G, in the human TRIM5α (huTRIM5α) domain responsible for retroviral capsid recognition leads to efficient HIV-1 restriction. Using a DNA transfection-based CRISPR-Cas9 genome editing protocol, we successfully mutated *TRIM5* to its HIV-1-restrictive version by homology-directed repair (HDR) in HEK293T cells. Nine clones bearing at least one HDR-edited *TRIM5* allele containing both mutations were isolated (5.6% overall efficiency), whereas another one contained only the R332G mutation. Of concern, several of these HDR-edited clones contained on-target undesired mutations, and none had all the alleles corrected. We observed a lack of HIV-1 restriction in the cell clones generated, even when cells were stimulated with IFN-I prior to infection. This, however, was partly explained by the unexpectedly low potential for TRIM5α-mediated restriction activity in this cell line. Our study demonstrates the feasibility of editing the TRIM5 gene to in human cells and identifies the main challenges to be addressed in order to use this approach to confer protection from HIV-1.

## Introduction

Viruses are obligate parasites whose success at infecting a host cell typically requires evasion from antiviral factors. In mammals, many cellular antiviral factors that can potentially interfere with the progression of viral infections have been identified. These factors can often act without external stimulation, but their expression and activity are enhanced by cytokines such as type I interferons (IFN-I) [1]. IFN-I cytokines are a multigene family of small peptides that include IFN-α, IFN-β, IFN-ε, IFN-κ, IFN-ω, IFN-δ and IFN-τ in humans [2]. Whereas these various IFN-I species all interact with the same receptor, they differ in IFN-stimulated gene (ISG)-, pathogen-and cell type-specificity [3].

Among the ISGs relevant to retroviruses, the family of viruses to which HIV-1 belongs, is *TRIM5*, which encodes the cytoplasmic protein TRIM5α [4]. In humans, *TRIM5* is transcribed into 5 isoforms, among which only TRIM5α possesses antiviral activity [5]. At its C-terminus, a domain called SPRY (PRYSPRY, B30.2) determines the retrovirus targeting specificity. This domain comprises hyper-variable loops that directly interact with the N-terminal domain of capsid proteins early after entry of the retrovirus into the host cell membrane [6]. When such interactions occurs, the retrovirus is inhibited (“restricted”) through mechanisms that include destabilization of the capsid core [7], proteasomal degradation of some core components [8] and sequestration of the viral particle in TRIM5α cytoplasmic bodies [9]. huTRIM5α generally has little-to-no activity against HIV-1, but efficiently inhibits the infectivity of the nonhuman gammaretrovirus “N-tropic” murine leukemia virus (N-MLV) as well as the nonhuman lentivirus equine infectious anemia virus (EIAV) [10]. Those two viruses are typically inhibited ∼10-fold (EIAV) and ∼100-fold (N-MLV) by endogenous huTRIM5α, with some variation depending on the cellular context.

Several groups, including ours, have attempted to harness the antiviral power of TRIM5α in order to interfere with HIV-1. This virus is efficiently restricted (∼100-fold) by some orthologs of TRIM5α found in Old World monkeys such as the Rhesus macaque TRIM5α (rhTRIM5α) [4]. However, significant sequence variation between the human and macaque orthologs preclude the possibility of using the latter one in gene therapy approaches, as this would increase the risk to elicit an immune response against the transgene in patients. Thus, all the studies have consisted in over-expressing versions of huTRIM5α designed to target HIV-1 through modifications in the SPRY domain. Some of the TRIM5α variants used were chimeric products containing small regions of rhTRIM5α in the SPRY [11,12]. Other teams mapped with further precision the HIV-1 restriction determinants in rhTRIM5α that were absent in huTRIM5α, leading to the discovery that mutating the Arg332 residue in huTRIM5α was sufficient to inhibit HIV-1. Although initial observations [13,14] raised the hope that single mutations at this position might inhibit HIV-1 as efficiently as rhTRIM5α did, later work made it clear that this was not the case [15].

Our laboratory explored a different approach: generating libraries of TRIM5α SPRY mutants then applying a functional screen to isolate mutants that conferred HIV-1 restriction [16,17]. These studies identified mutations at Arg335 inhibiting HIV-1 by 5-to 10-fold. When we combined a mutation at Arg335 (R335G) with one at Arg332 (R332G), we obtained restriction levels that were higher than with either of the single mutants [16,17]. Although not quite as restrictive as rhTRIM5α, R332G-R335G huTRIM5α efficiently inhibited the propagation of a highly pathogenic strain of HIV-1, and cells expressing the transgene had a survival advantage over unmodified cells [15].

Although R332G-R335G huTRIM5α is considered a prime candidate in HIV-1 gene therapy approaches to inhibit HIV-1, using lentiviral vectors to overexpress it in human cells is not without caveats. Indeed, the physiological effects of TRIM5α overexpression *in vivo* are not clear, considering that it is involved in innate immune responses [18,19] and possibly in autophagy [20]. In addition, the genotoxicity of lentiviral vectors integrating in the human genome is still poorly predictable. In the longer term, it would thus be desirable to be able to introduce mutations in the endogenous human *TRIM5* by genome editing. Here we describe the use of Clustered Regularly Interspaced Short Palindromic Repeats with Cas9 (CRISPR-Cas9) and of a single-stranded homology-directed repair (HDR) donor DNA to successfully mutate Arg332 and Arg335 in a human cell line.

## Results

### Strategy for the mutagenesis of *TRIM5* by HDR

Human *TRIM5* is found on chromosome 11p15.4 and the region of the gene encoding the SPRY domain is present in exon 8 (Fig 1A). We searched for DNA loci close to the codons for Arg332 and Arg335 that would be potential targets for CRISPR-Cas9-mediated double-strand cleavage. CRISPR guide RNAs (gRNAs) were designed for the 3 potential target sites that were nearest to the two codons to be mutated (Fig 1B). CRISPR plasmids expressing Cas9 along with one of the designed gRNAs were transfected in human embryonic kidney 293T cells (HEK293T), and a Surveyor assay was performed to test the capacity of the three gRNAs to target the endogenous TRIM5 gene. Results showed that all three gRNAs were competent (Fig 1B). Because gRNA1 induces a cleavage that is closest to the targeted codons (right before the first nt of Arg332), the rest of the project was carried out with this gRNA. The HDR donor DNA consisted of a single-stranded oligodeoxynucleotide (ssODN) that was 200 nt long and antisense relative to the gRNA, in keeping with published methods [21,22]. The central section of this ssODN containing the mutations introduced is shown in Fig 1C (depicted in the same orientation as the TRIM5 mRNA for clarity purposes). In addition to the mutations substituting arginine residues into glycine at positions 332 and 335, we included 4 silent mutations in the region recognized by the gRNA and 1 more silent mutation in the protospacer adjacent motif (PAM), amounting to a total of 7 substitutions expected to prevent the cleavage of the donor DNA by Cas9. One of the mutations also created a HaeIII cut site for convenient downstream screening of the cell clones obtained.

**Fig 1.**
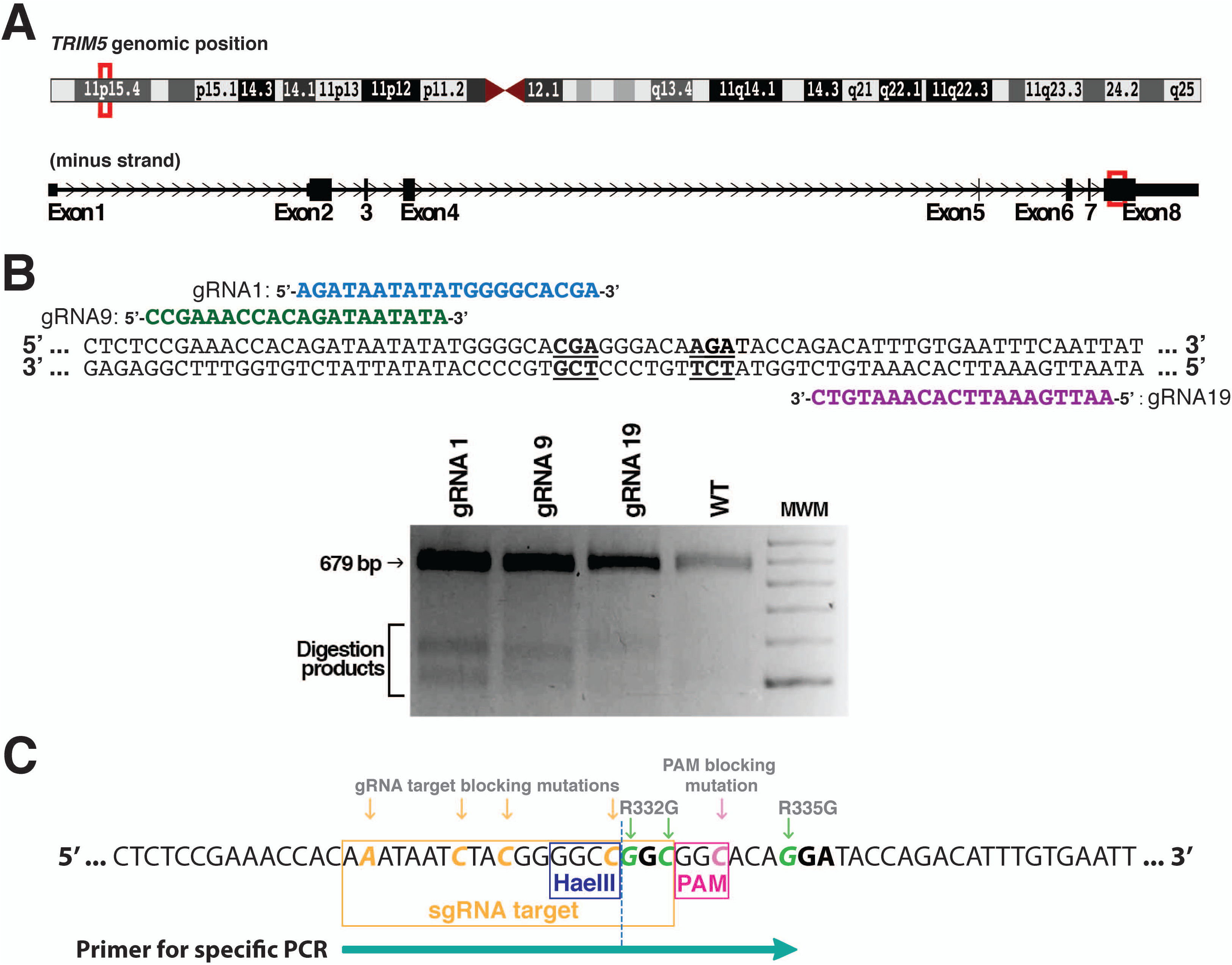
Design of the gRNA and donor ssODN for the HDR-mediated editing of *TRIM5*. (A) *TRIM5* localization on chromosome 11 (top), and Arg332-Arg335 localization in exon 8 of the gene (bottom). (B) Top panel: position of the three gRNAs (gRNA1, 9 and 19) designed to target the Arg332-Arg335 region. The two arginine codons are underlined and in bold. Bottom panel: Surveyor assay following the transfection of HEK293T cells with CRISPR-Cas9 plasmids expressing one of the three gRNAs. WT DNA from nontransfected cells was used as a control. (C) HDR donor DNA mutagenesis strategy. 8 substitutions were present, including three nonsilent substitutions to mutate Arg332 and Arg335 into Gly (green), one silent mutation to disrupt the PAM sequence (pink), and four silent mutations in the sequence targeted by gRNA1 (orange). The HaeIII restriction site created as a result of one of the silent substitutions is indicated, as is the position of the primer used in specific PCR screening.

### Isolation of *TRIM5*-edited HEK293T clones

We transiently transfected a plasmid (pX459) expressing Cas9 and the gRNA1 into HEK293T cells, along with the ssODN. Single-cell clones were then isolated by limiting dilution. We screened 161 clones at random for the presence of HDR-modified alleles by specifically amplifying the mutated TRIM5 sequence using a primer whose sequence is indicated in Fig 1C. As shown in Fig S1, 14 clones showed a positive signal (of varying intensity) in this assay. One of the clones, F2X, yielded a band whose size seemed bigger compared to another positive clone (C8) analyzed on the same gel. All these clones (minus A12, which did not survive) were re-analyzed using the same specific PCR assay and also using a second assay in which the targeted region is amplified and then digested by HaeIII, which cuts at a site created by one of the silent mutations (see Fig 1C). In the latter assay, amplification was done using primers that bind outside the 200 nts corresponding to the donor ssODN in order to insure that the ssODN was not inadvertently detected. Figure 2 shows a positive signal for 10 of these clones in both assays. The three remaining clones were negative in both assays.

**Fig 2.**
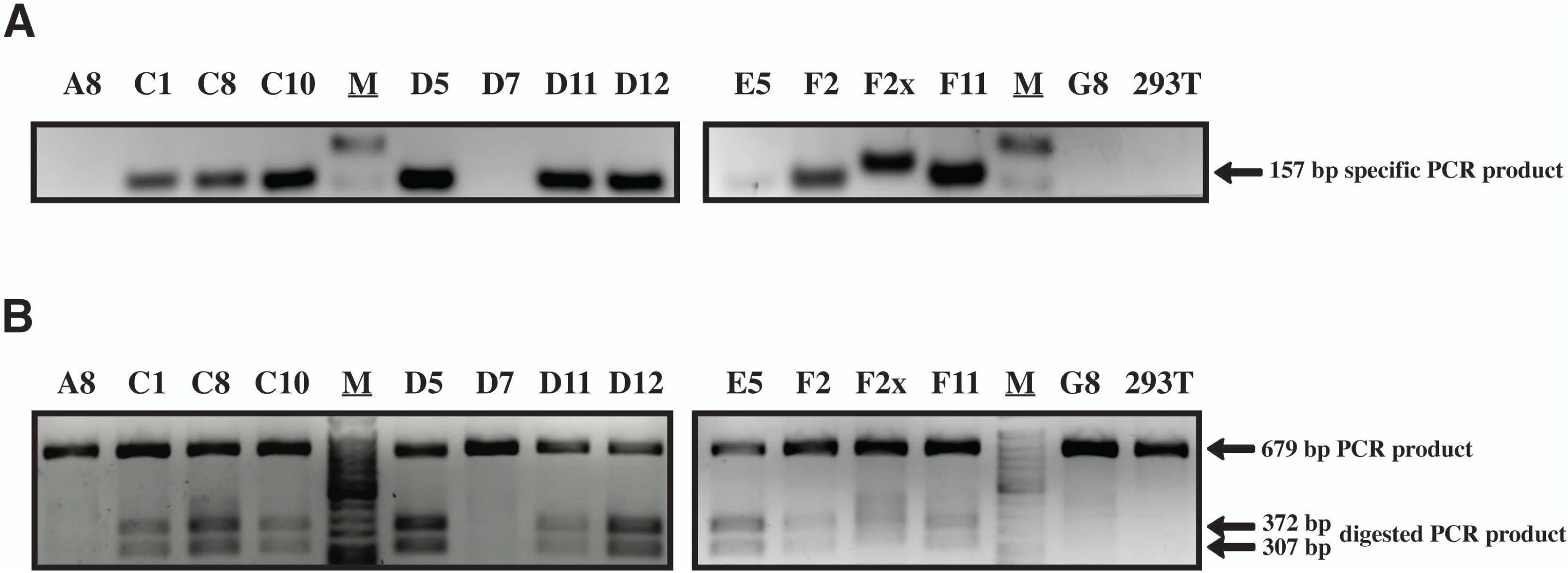
Identification of HDR-edited clones. Following the isolation of HEK293T clones, HDR-edited clones were identified by a dual PCR screen. The figure shows the analysis of 13 clones that passed a pre-screen step (see Supporting Information). (A) PCR using a primer specific for the mutated *TRIM5*. Untransfected HEK293T cells were used as a control. M, molecular weight marker. (B) Non-specific PCR of the targeted region followed by HaeIII digestion. The expected sizes of the digested PCR products are shown on the right. The full-length gels are available on the FigShare public repository (see “Availability of data” section).

### Genotype analysis

The 10 clones showing indications of HDR-mediated editing were then subjected to PCR using the same primers that were also utilized in the HaeIII screen described above. PCR products were analyzed by deep sequencing, and a color-coded alignment of the results is shown in Fig 3. The HEK293 cells and their derivatives are pseudotriploid [23] and accordingly, we found that two clones had two *TRIM5* alleles and seven were triploid. F2X seemed to possess 6 *TRIM5* alleles, a finding that is discussed below. For each clone, both desired mutations at Arg332 and Arg335 were present on one allele or more, with the exception of F2 which only had the Arg332 mutation. However, only one clone (D11) contained an allele with all 8 substitutions present. The HDR-generated alleles in the other cell clones generally contained the expected mutations in the PAM-proximal side of the cleavage site, with the exception of F2 which lacked the A-to-G mutation required to introduce the R335G change. All HDR-generated alleles had the A-to-C silent mutation creating the HaeIII restriction site at the first nucleotide upstream of the cleavage site on the PAM-distal side. This is consistent with the fact that all clones that were found to be positive in the specific PCR screen were also positive in the HaeIII assay (Fig 2). Strikingly, for the three other substitution mutations in the PAM-distal region, only one clone (D11) had all of them whereas another one (F11) had only one. It would be tempting to conclude that HDR is biased so that mutations were more likely to be incorporated in the PAM-proximal side of the cut, and other teams have reported such imbalances in the conversion rate [24]. On the other hand, our specific PCR screen requires a successful amplification using a primer whose 3’ half is complementary to the PAM-proximal region (Fig 1C), thus creating a bias toward the detection of mutated DNA containing the expected mutations in this PAM-proximal region.

**Fig 3.**
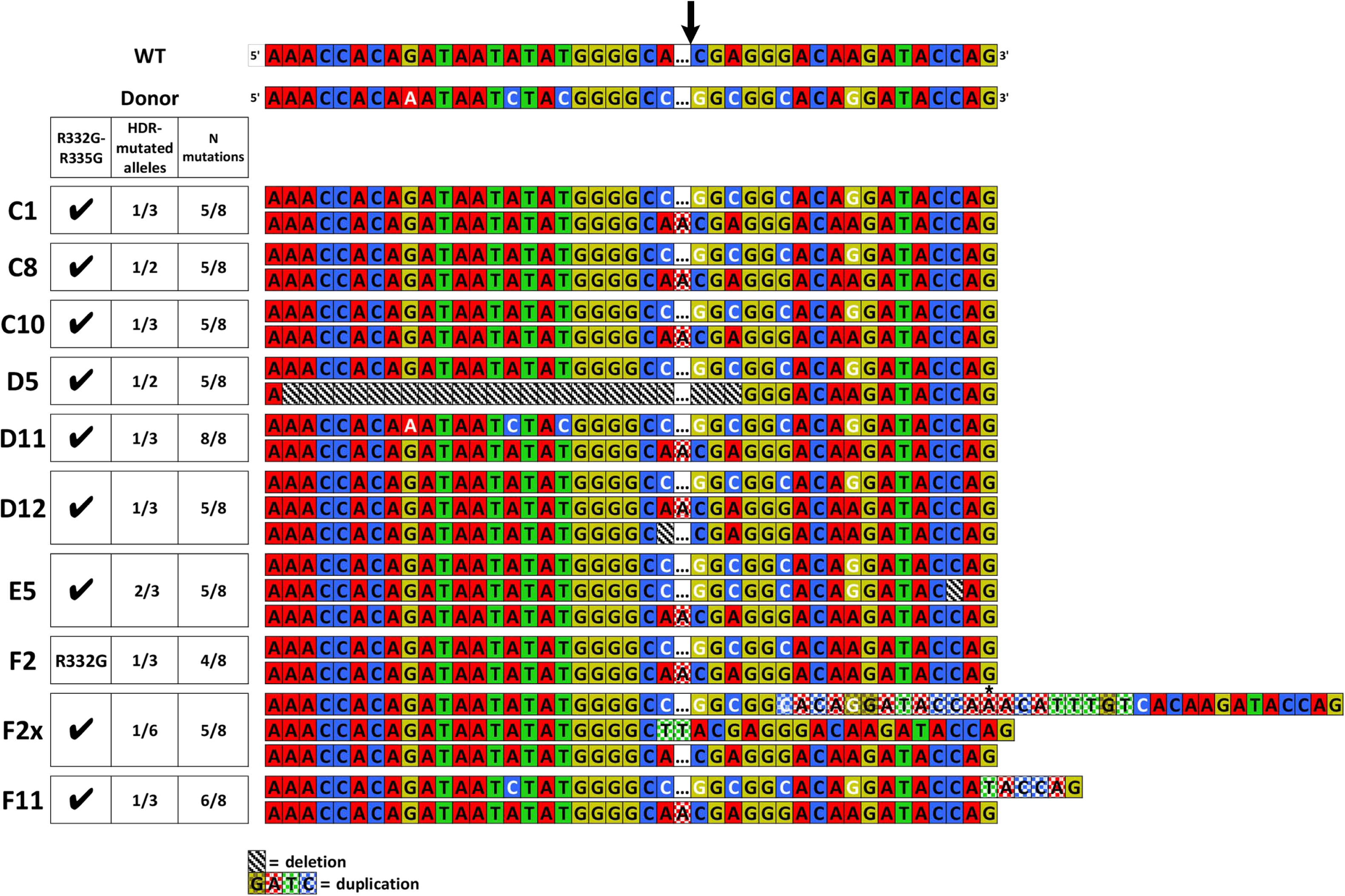
Deep sequencing analysis of *TRIM5* editing in 10 screened clones. The ∼200-nt HDR-targeted *TRIM5* region was amplified by PCR and the PCR products were then analyzed by Illumina MiSeq sequencing. The alignment shown includes the targeted locus for each allele of the 10 clones, in comparison with the WT sequence and with the expected HDR-mutated sequence (top 2 lines). Expected substitutions are shown in white. The star indicates the position of an unexpected substitution within a duplicated region in one allele of F2X. The color code for duplications/insertions and for deletions is explained at the bottom of the alignments. The Cas9 cleavage site on the WT sequence is shown at the top. On the left is a table summarizing the results obtained for each clone: presence of the two therapeutic mutations R332G/R335G, proportion of *TRIM5* alleles modified by HDR and the proportion of the expected substitution mutations in the HDR-edited alleles. Note that only one clone (D11) has an allele containing all the desired mutations and that most of the non-HDR-edited alleles contain indels at the cleavage site.

Some HDR-generated alleles showed additional, unexpected mutations. In E5, one of the two HDR alleles had a one-nt deletion deletion 16 nts from the cleavage site in the PAM-proximal region. In F11, the HDR allele had a 5 nt (TACCA) duplication in the same region. F2X showed an intriguing genotype: firstly, we found that the HDR allele was present in 17 % (1:6) of the amplicons, whereas 33 % (2:6) were wild-type (WT) and 50% (3:6) contained a TT insertion. HEK293 cells are known to be prone to chromosomal translocations leading to a high level of variation in copy numbers [25], which might explain our findings. The HDR-generated allele in F2X had a surprising structure, with a 21 nt duplication consistent with the slower-migrating bands in Fig 2 and in Fig S1. The repeated sequence that was closest to the cleavage site had the expected mutations in the PAM and at Arg335 and also contained an additional substitution (G-to-A) that is not present in the donor ssODN, whereas the second repeat of this sequence only had the G-to-C mutation in the PAM. These HDR-generated alleles that also contained unexpected insertions/deletions (indels) are unlikely to encode functional TRIM5α, due to the frameshifts leading to premature termination (E5, F11) or due to the insertion of 7 aminoacids at a region crucial for capsid binding (F2X). The rest of the HDR-generated alleles (in C1, C8, C10, D5, D12 and one of the two HDR alleles in E5), however, may potentially encode proteins that efficiently target HIV-1.

Examination of the *TRIM5* alleles not modified by HDR in these cell clones revealed that they showed clear signs of NHEJ-induced mutations, i.e. indels at the cleavage site. 8 alleles had an A inserted at the cleavage site, whereas one of the F2X alleles had a TT inserted at the same locus. The A insertion was so prevalent that in half of the clones (C1, C10, D11, F2, F11), 2 of the 3 *TRIM5* copies were mutated by NHEJ leading to this particular mutation. Although the nature of NHEJ-directed mutations is known to vary widely depending on the gRNA used [26], such +1 insertions have been described to be prevalent as a result of Cas9 editing [27,28]. One D5 allele had a larger, 27 nt-long deletion whereas one D12 allele had a single deletion at the cleavage site. Therefore, and with the exception of F2X, our data strongly suggest that in all cell clones in which one or two of the *TRIM5* alleles were mutated by HDR, the remaining alleles were mutated by NHEJ. This is consistent with findings published by others [21]. Furthermore, these NHEJ-generated *TRIM5* alleles are all expected to encode non-functional TRIM5α due to missense mutations in the SPRY domain and premature termination.

### Lack of HIV-1 restriction activity in the *TRIM5*-edited cells

We challenged the 10 cell clones with HIV-1_NL-GFP_, an HIV-1-derived vector that was previously used extensively to study TRIM5α [29]. Cells were infected with several different amounts of the GFP-encoding vector, and the % of GFP-positive cells was calculated by FACS as a measurement of infectivity (Fig 4A). Compared with the parental HEK293T cells, some of the clones showed a slightly increased permissiveness (up to 3-fold) whereas others were slightly less permissive to the infection (up to 2-fold). Because of the expected cell-to-cell variation in susceptibility to retroviral infection, irrespective of TRIM5α, we challenged the same cell populations with other retroviral vectors, derived from the nonhuman primate lentivirus SIVmac239, the equine lentivirus EIAV, as well as the huTRIM5α-sensitive murine oncoretrovirus N-MLV and its huTRIM5α-insensitive counterpart B-MLV. In previous studies, we found that compared to the WT huTRIM5α, the R332G-R335G mutant restricted SIVmac and N-MLV at slightly higher levels, whereas EIAV and B-MLV were not affected [16]. Therefore, if HIV-1 was specifically restricted by a TRIM5α mutant in a given cell clone, we would expect the EIAV and B-MLV vectors to be restriction-negative controls. However, the infectivity profiles obtained in the various cell populations were similar. In particular, C10 was the cell clone least permissive to HIV-1_NL-GFP_ (Fig 4A), but it was also poorly permissive toward infection by the 4 other retroviral vectors (Fig 4B-E). Likewise, C1 was relatively more susceptible to infection by HIV-1_NL-GFP_, but these cells were also more permissive to infection with the other 4 retroviral vectors. Of interest, however, was the fact that N-MLV_GFP_ was less infectious in the parental cells compared with most of the cell clones, hinting at slightly higher relative levels of restriction in the parental cells for this virus. In conclusion, the modest variations in HIV-1 infectivity observed between cell clones result from cellular factors other than TRIM5α, and instead are probably due to variations in the expression levels of “positive” factors such as the receptor for the viral vector envelope used here.

**Fig 4.**
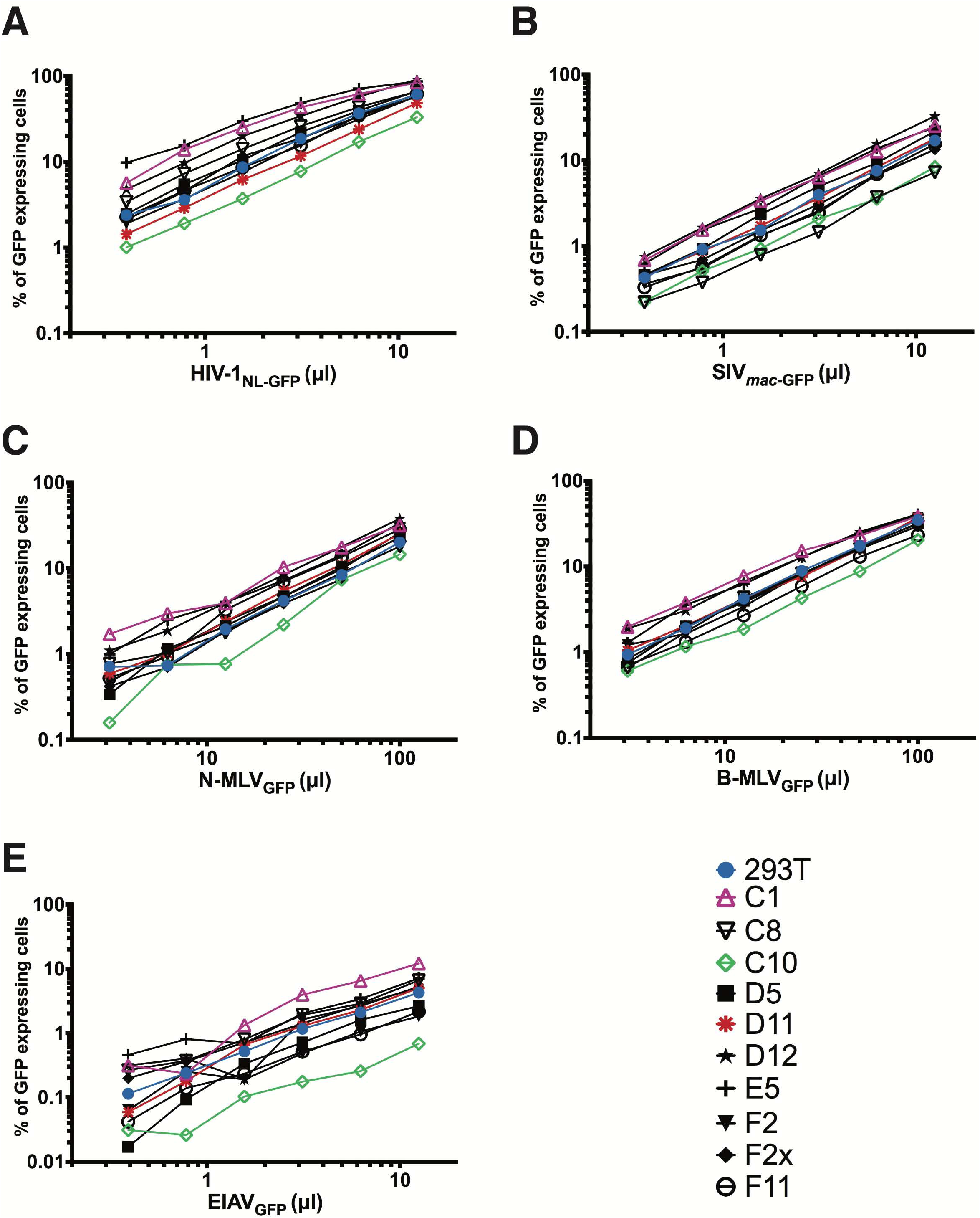
Retrovirus restriction ability of *TRIM5*-edited clones. The 10 HDR-edited clones were challenged with increasing doses of GFP-expressing retroviral vectors based on HIV-1_NL4-3_ (A), SIV_mac239_ (B), N-MLV (C), B-MLV (D) or EIAV (E). Non-transfected HEK293T cells were used as a control. Infected cells were quantified by FACS for GFP expression.

### Interferon treatment does not induce HIV-1 restriction in *TRIM5*-edited cells

In the infection experiment described above, we were surprised to find that the N-MLV vector was more infectious than is usually observed in human cells. Indeed, N-MLV is usually restricted ∼100-fold in human cells such as HeLa cells [30], whereas the restriction level was closer to 3-fold in our HEK293T cells. It is possible that TRIM5α expression levels are lower in this particular cell line. TRIM5α transcription is stimulated by IFN-I, especially IFN-β [31]. Therefore, we reasoned that IFN-I treatment might reveal a restriction activity against HIV-1 and N-MLV by boosting TRIM5α levels. To test this possibility, we challenged the parental HEK293T cells as well as each of the 10 clones with the HIV-1, SIVmac, N-MLV and B-MLV vectors and in the absence or presence of IFN-α, IFN-β and IFN-ω (Fig 5). In the parental cells, we observed that treatment with IFN-α and IFN-β, but not IFN-ω, slightly inhibited the infectivity of the HIV-1 vector (Fig 5A) but had a relatively bigger effect (2-fold) on the N-MLV vector (Fig 5C). IFN-β had a very small inhibitory effect on B-MLV, whereas the two other IFN-I species did not affect the infectivity of this virus (Fig 5D). These results are consistent with IFN-α and IFN-β (but not IFN-ω) enhancing TRIM5α expression, hence leading to a specific increase in N-MLV restriction. In the absence of IFN-I, the permissiveness of the cell populations to HIV-1_NL-GFP_ infection varied within a ∼4-fold window (Fig 5A), similar to what we observed in the experiment shown in Fig 4. IFN-I had little-to-no effect on the infectivity of HIV-1_NL-GFP_ in the 10 cell clones analyzed, implying that the treatment did not induce restriction. Interestingly, N-MLV_GFP_ infectivity was similarly not decreased by IFN-I treatment in the 10 edited cell clones except for D5, and in the presence of IFN-α or IFN-β, N-MLV_GFP_ was more infectious in all of the clones than it was in the treated parental cells. These observations suggest that N-MLV restriction was lost in the cell clones due to the mutations introduced in *TRIM5*.

**Fig 5.**
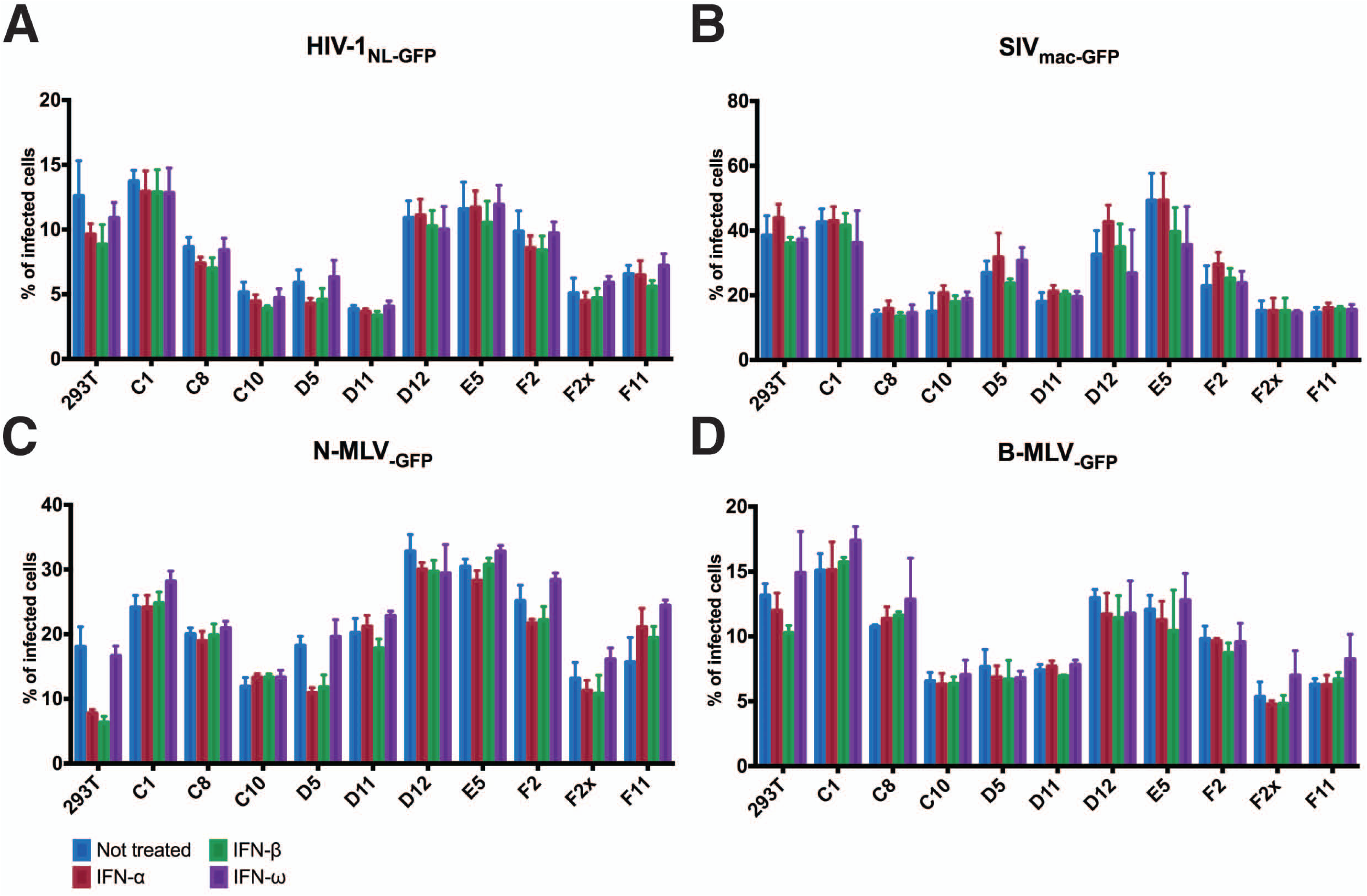
Retrovirus restriction ability following IFN-I treatment. HDR-edited clones and the nontransfected WT control cells were treated with IFN-α, IFN-β or IFN-ω for 16h prior to a single-dose infection with HIV-1_NL-GFP_ (A), SIV_mac-GFP_ (B), N-MLV_GFP_ (C) and B-MLV_GFP_ (D). The percentage of infected cells was determined by FACS.

### HEK293T cells are suboptimal for TRIM5α-mediated restriction

Since we had unexpectedly observed that N-MLV was not efficiently restricted by the WT endogenous *TRIM5* in the HEK293T cells used here, we scrutinized its sequence more closely. A published study showed that *TRIM5* in this cell line carries a SNP resulting in the R136Q mutation [25]. The effects of this mutation on retroviral restriction are unclear, with some authors linking it to an increase in susceptibility toward HIV-1 whereas others find the opposite [32]. The presence of this SNP might explain the weak restriction levels observed in this study. In addition, HEK293T cells might provide an inadequate environment for TRIM5α to restrict retroviruses, irrespective of the R136Q mutation in *TRIM5*. To test this possibility, we stably transduced R332G-R335G huTRIM5α into 293s. In other cell lines tested, transduction of this double mutant leads to a ∼20-fold decrease in HIV-1 infectivity [16,17,29]. In HEK293T cells, however, we observed restriction levels that were closer to 3-fold (Fig S2). We conclude that the potential for TRIM5α-mediated restriction is abnormally low in the HEK293T cell line used here.

## Discussion

To our knowledge, this is the first attempt at using genome editing to mutate a restriction factor with the aim of conferring an innate antiviral function in human cells. Although we did not detect an antiviral activity following successful *TRIM5* editing events, we identified possible reasons for this: (i) the presence of the R136Q mutation in *TRIM5* in this cell line; (ii) the fact that this specific cell line provided a suboptimal environment for retroviral restriction by TRIM5α; and finally, (iii) the co-presence of indel-containing *TRIM5* alleles in all cell clones in which an allele had been modified by HDR. It is predicted that the SPRY-truncated TRIM5α proteins resulting from the presence of indels will interact with the full-length TRIM5α and will interfere with its targeting of incoming retroviruses, similar to the activity of natural, shorter TRIM5 isoforms [5,33]. We did not investigate any further the reasons behind the lack of restriction in this particular cell line, as we are now focusing on editing *TRIM5* in lymphoid and myeloid cells. However, future research will need to address the difficulty of achieving bi-allelic HDR-mediated editing. Recent technological advances, including the development of a marker-free system to enrich cells in which HDR occurred [34], are likely to enhance editing efficiency as well as bi-allelic editing. Also affecting the therapeutic potential of *TRIM5* editing is the occurrence of unwanted on-target indels, in 3 out of 9 alleles bearing the two therapeutic mutations in this study. Therefore, future studies will also need to identify the determinants of fidelity in HDR repair in order to minimize the incidence of such indels. Despite the absence of an antiviral effect, this study paves the way and identifies the pitfalls toward the goal of efficient, bi-allelic, scarless *TRIM5* editing in order to confer HIV-1 resistance in human cells.

## Methods

### Cells and *TRIM5* genotyping

HEK293T cells were maintained in Dulbecco’s modified Eagle’s medium (DMEM; HyClone). All culture media were supplemented with 10% fetal bovine serum (FBS) and penicillin/streptomycin (HyClone). To analyze the sequence of the targeted genomic region, cellular DNA was prepared using the Bioline genomic DNA kit (London, UK), and the *TRIM5* region encompassing the targeted locus was PCR-amplified using primers T5a_Surveyor_fwd (5’GTCCGACGCTACTGGGGTAAG) and T5a_Surveyor_rev (5’ATAATCACAGAGAGGGGCACA). The PCR product was Sanger-sequenced using the same primers. We found no variation in this region compared to the consensus sequence (NCBI Gene ID: 85363).

### Design of gRNAs and Surveyor assay

The lentiviral expression vector pLentiCRISPRv2 (pLCv2) was a gift from Feng Zhang (Addgene plasmid # 52961) [35]. Three gRNAs targeting *TRIM5* were designed using the Zhang lab online software available at crispr.mit.edu. The sequences targeted are 5’AGATAATATATGGGGCACGA.(gRNA1),.5’CCGAAACCACAGATAATATA (gRNA9) and 5’AATTGAAATTCACAAATGTC (gRNA19). The ODNs needed for the generation of pLCv2-based constructs targeting those sequences were designed exactly as described in published protocols [26,35]. Sense/antisense pairs of primers were annealed and cloned into pLCv2 cut with BsmBI. To evaluate the capacity of the constructed plasmids to induce on-target indels in *TRIM5*, a surveyor nuclease assay was performed. HEK293T cells were transfected with either pLCv2-gRNA1, -gRNA9 or -gRNA19 using polyethyleneimine [36]. 3 d later, the genomic DNA was extracted from the transfected cells using the Bioline genomic DNA kit. The targeted *TRIM5* region was PCR-amplified using primers T5a_Surveyor_fwd and T5a_Surveyor_rev. PCR amplicons were heat-denatured at 95°C, and re-annealed by slow cooling to promote formation of dsDNA heteroduplexes. The heteroduplexes were then cleaved by Surveyor nuclease S provided as part of the Transgenomic Surveyor mutation detection kit (Integrated DNA Technologies, Coralville, IA), according to the manufacturer’s instructions. Digestion products were visualized by agarose gel electrophoresis.

### Design of the HDR donor DNA and *TRIM5* editing

The following *TRIM5* minus strand-derived HDR DNA was synthesized by Integrated DNA Technologies: 5’CGTCTACCTCCCAGTAATGTTTCCCTGATGTGATACTTTGAGAGCCCAGGATGCCA.GTACAATAATTGAAATTCACAAATGTCTGGTATCCTGTGCCGCCGGCCCCGTAGATT.ATTTGTGGTTTCGGAGAGCTCACTTGTCTCTTATCTTCAGAAATGACAGCACATGAAATGTTGTTTGGAGCCACTGTCACATCAACT. Residues mutated compared to the WT *TRIM5* sequence are underlined. pX459-gRNA1 was constructed in a manner similar to pLCv2-gRNA1, by ligating the corresponding annealed gRNA1 ODN duplex into pX459 (pSpCas9(BB)-2A-Puro; Addgene #62988) [37] digested with BbsI. HEK293T cells were plated in 6-well plates at 2.7 × 10^5^ per well and transfected the next day using polyethyleneimine, with 2.5 μg of pX459-gRNA1 together with 5 μl of the HDR DNA prepared at 20 μM. When cells reached confluence, they were trypsinized and plated at 0.5 cell per well in 96-well plates, using conditioned medium. To screen the colonies for HDR-mediated *TRIM5* editing, part of the cells were lyzed in the DirectPCR Lysis reagent (Viagen Biotech, Los Angeles, USA) diluted 1:1 in proteinase K-containing water as recommended by the manufacturer. Lysis was allowed to proceed overnight at 55°C followed by heating at 85°C for 90 min to deactivate proteinase K. For the specific PCR-based screening, 5 μl of the lysed cells were subjected to PCR using primers T5a_mut_fwd (5’-AAATAATCTACGGGGCCGGCGGCACAG) and T5a_qPCR_rev (5’-CCAGCACATACCCCCAGGAT). PCR was performed for 30 cycles using the following conditions: 30 sec at 94°C, 30 sec at 61.5°C, 30 sec at 68°C. The 157-bp expected PCR product was resolved on agarose gels. For the HaeIII-based screening, lysed cells were subjected to PCR using primers T5a_Surveyor_fwd and T5a_Surveyor_rev, similar to the Surveyor assay. 10 μl of the PCR product were digested by HaeIII for 60 min at 37°C. The reaction products were analyzed using agarose gels in order to reveal the 307-bp and 372-bp bands corresponding to digested products. For MiSeq sequencing, cellular DNA was submitted to PCR using the following ODNs, which bind to DNA sequences located 10 nt outside the 200 nt-long region corresponding to the HDR: huTR5aGG_seq_FOR, 5’ACACTGACGACATGGTTCTACAATCCCTTAGCTGACCTGTTA, and.huTR5aGG_seq_REV,5’TACGGTAGCAGAGACTTGGTCTCCCCCAGGATCCAAGCAGTT. The underlined sequences are barcodes. MiSeq sequencing results were analyzed using the online tool Integrative Genomics Viewer (http://software.broadinstitute.org/software/igv/).

### Retroviral vectors production and viral challenges

To generate the HEK293T cells stably expressing WT and R332G-R335G huTRIM5α, cells were transduced with the corresponding pMIP-huTRIM5α vectors followed by puromycin selection as described previously [16]. To produce GFP-expressing retroviral vectors, HEK293T cells were seeded in 10 cm culture dishes and transiently co-transfected with the following plasmids: pMD-G, pCNCG and pCIG3-B or pCIG3-N to produce B-MLV_GFP_ and N-MLV_GFP_, respectively; pMD-G and pHIV-1_NL-GFP_ to produce HIV-1_NL-GFP_; pMD-G and pSIV_mac239-GFP_ to produce SIV_mac-GFP_; or pONY3.1, pONY8.0 and pMD-G to produce EIAV_GFP_ (see [38,39] and references therein). For retroviral challenges, cells were seeded into 96-well plates at 10,000 cells per well and infected the following day with multiple doses of the GFP-expressing retroviral vectors. Cells were trypsinized at 2 d post-infection and fixed in 2.5% formaldehyde (Fisher Scientific, MA, USA). The percentage of GFP-positive cells was then determined by analyzing 1 × 10^4^ cells on a FC500 MPL cytometer (Beckman Coulter, CA, USA) using the CXP Software (Beckman Coulter). For infections done in presence of IFN-I, recombinant human IFN-α, IFN-β or IFN-ω (PeproTech, Rocky Hill, NJ) was added to cell cultures 16 h prior to infection and at a final concentration of 10 ng/ml.

### Availability of materials and data

The datasets generated in the course of this study are available in the FigShare repository, https://figshare.com/projects/Dufour_et_al_2017/24445. Biological materials are available for sharing.

## Acknowledgements

We are grateful to Feng Zhang (Broad Institute, Cambridge, USA) for sharing reagents. We thank Pierre Lepage and Frédérick Robidoux (Genome Quebec Innovation Center, Montréal) for help with MiSeq data generation and analysis.

## SUPPORTING INFORMATION CAPTIONS

**Fig S1. Pre-screening of 161 isolated clones by specific PCR.** Following 20 days of growth, 161 isolated HEK293T clones were screened for HDR-edited TRIM5 gene by mutation-specific PCR. 14 clones that passed this pre-screen step are indicated by their names. MWM, molecular weight marker.

**Fig S2. Levels of HIV-1 restriction in HEK293T cells transduced with R332G-R335G huTRIM5α.** HEK293T cells were retrovirally transduced with WT huTRIM5α, R332G-R335G huTRIM5α or with the “empty” vector as indicated. Untransduced cells were eliminated, and the cell populations were then challenged with increasing amounts of the HIV-1_NL-GFP_ vector. The percentage of cells expressing GFP was then determined by FACS.

